# A novel pathway controlling cambium initiation and - activity via cytokinin biosynthesis in Arabidopsis

**DOI:** 10.1101/2020.06.19.162297

**Authors:** Arezoo Rahimi, Omid Karami, Angga Dwituti Lestari, Dongbo Shi, Thomas Greb, Remko Offringa

**Affiliations:** Plant Developmental Genetics, Institute of Biology Leiden, Leiden University, Sylviusweg 72, 2333 BE Leiden, The Netherlands; Department of Developmental Physiology, Centre for Organismal Studies (COS), Heidelberg University, 69120 Heidelberg, Germany

**Keywords:** Secondary growth, *AHL15*, *SOC1*, *FUL*, cytokinin, Arabidopsis

## Abstract

Plant secondary growth, also referred to as wood formation, includes the production of secondary xylem, which is derived from meristematic cambium cells embedded in vascular tissues. Despite the importance of secondary xylem in plant growth and wood formation, the molecular mechanism of secondary growth is not yet well understood. Here we identified an important role for the *Arabidopsis thaliana* (Arabidopsis) *AT-HOOK MOTIF CONTAINING NUCLEAR LOCALIZED 15 (AHL15)* gene, encoding for a putative transcriptional regulator, in controlling vascular cambium activity and secondary xylem formation. Secondary xylem development was significantly reduced in inflorescence stems of the Arabidopsis *ahl15* loss-of-function mutant, whereas *AHL15* overexpression led to extensive secondary xylem formation. *AHL15* expression under a vascular meristem-specific promoter also enhanced the amount of interfascicular secondary xylem. Moreover, *AHL15* appeared to be required for the enhanced secondary xylem formation in the Arabidopsis double loss-of-function mutant of the *SUPPRESSOR OF OVEREXPRESSION OF CO 1* (*SOC1*) and *FRUITFULL* (*FUL*) genes. A well-known central regulator of cambial activity is the plant hormone cytokinin. We showed that the expression of two cytokinin biosynthesis genes (*ISOPENTENYL TRANSERASE (IPT) 3* and *7*) is decreased in *ahl15* loss-of-function mutant stems, whereas the secondary xylem deficiency in these mutant stems can be resorted by cambium-specific expression of the *Agrobacterium tumefaciens IPT* gene, indicating that *AHL15* acts through the cytokinin pathway. These findings support a model whereby *AHL15* acts as a central factor inducing vascular cambium activity downstream of *SOC1* and *FUL* and upstream of *IPT3*, *IPT7* and *LOG4, LOG5* governing the rate of secondary xylem formation in Arabidopsis inflorescence stems.

## Introduction

Throughout their lifespan, plants can dynamically change their growth and development in response to environmental signals, and this allows them to adapt and survive adverse conditions. In many flowering plants, but especially in woody plant species, stems show two distinct developmental growth processes. The increase in stem length and the establishment of the primary vascular meristem that is arranged into vascular bundles in young stems are known as primary growth. Each bundle contains a primary vascular meristem, or a procambium, that generates primary xylem towards the center and primary phloem towards the periphery of the stem (Fischer et al., 2019). Later in development, when primary stem growth is completed, the procambium and its neighbouring interfascicular parenchyma cells (between the vascular bundles) differentiate into the cambial meristem, eventually forming a continuous cylinder of stem cells in the primary plant stem. This then initiates the process of secondary growth, during which the cambial meristem continuously generates secondary xylem usually towards the inside of the stem, and secondary phloem towards the outside of the stem, resulting in radial stem expansion (Jouannet et al., 2015).

The vascular cambium is an important meristem, as it produces vascular tissues that transport water and nutrients throughout the plant body. Through secondary growth, it provides mechanical support for the plant body by generating lignified wood cells. The rate of cell division in stem cells of the vascular cambium determines the amount of secondary growth and, thus, whether and how much wood is formed. In view of the use of wood as building material and renewable energy source, there is an obvious demand for an understanding of the genetic mechanisms that control the cambium activity. The model plant *Arabidopsis thaliana* (Arabidopsis) is herbaceous, but it can undergo secondary growth in the hypocotyl, root, or stem depending on the growth conditions (Ragni and Greb, 2018; Fischer et al., 2019). Most of our current understanding of secondary growth comes from studies in Arabidopsis. Although great progress has been made over the past few years (insert references), the exact signals that initiate the formation of the cambium ring and regulate the amount of secondary growth, and thus differentiate between woody and non-woody species, are still unknown.

Previous studies have revealed a number of molecular factors and genetic components, including phytohormones and transcription factors, that are involved in the establishment and proliferation of cambium cells (Oh et al., 2003; Fischer et al., 2019). The TDIF-PXY-WOX signaling pathway is the best-studied signaling pathway that controls cambium activity. The peptide ligand CLE41/44/TRACHEARY ELEMENT DIFFERENTIATION INHIBITORY FACTOR (TDIF) is synthesized in the phloem and seems to diffuse through the apoplastic space to cambium cells, where it binds to its cognate receptor PHLOEM INTERCALATED WITH XYLEM (PXY). PXY subsequently promotes the proliferation of cambium cells by activating the expression of the cambium-specific *WUSCHELs-RELATED HOMEOBOX* genes *WOX4* and *WOX14* (Etchells and Turner, 2010; Hirakawa et al., 2010; Wang et al., 2019). The WOX4 transcription factor promote stem cell proliferation by interacting with the GRAS domain transcription factor HAIRY MERISTEM 4 (HAM4) (Zhou et al., 2015).

Cambium activity is also known to be regulated via a hormonal signaling network (Bhalerao et al., 2016; Immanen et al., 2016; Brackmann et al., 2018; Fischer et al., 2019). Cytokinin is a major plant hormone that is well-known to promote cell division in various meristems. In parallel to the TDIF-PXY-WOX pathway, cytokinin signaling has been highlighted as a major positive regulator of vascular cambium cell proliferation during secondary growth (El-Showk et al., 2013). ISOPENTENYL TRANSFERASE (IPT) enzymes are mostly responsible for cytokinin biosynthesis. The Arabidopsis *ipt1,3,5,7* quadruple mutant completely lacks cambial activity, but this can be rescued by treatment with exogenous cytokinin (Miyawaki et al., 2006; Matsumoto-Kitano et al., 2008). Moreover, the expression of a cytokinin catabolic gene in *Populus alba* (poplar) led to a remarkable decrease in cambial cell divisions and thinner trunks (Immanen et al., 2016). By contrast, increasing cytokinin levels in poplar stems by overexpressing *IPT7* under the wood-specific *LMX5* promoter strongly stimulated cambial cell division and biomass production (Immanen et al., 2016). In addition, it was shown that secondary growth in *Arabidopsis thaliana* (Arabidopsis) roots is dependent on the cytokinin-responsive transcription factor AINTEGUMENTA (ANT) and the D-type cyclin CYCD3;1, which are both expressed in the vascular cambium (Dewitte et al., 2007; Randall et al., 2015). Although a positive role for cytokinin signaling in cambial stem cell activity has been demonstrated, it is still largely unknown how cytokinin-signaling promotes cambial activity.

Here we identified a new role for the Arabidopsis *AT-HOOK MOTIF CONTAINING NUCLEAR LOCALIZED 15 (AHL15)* gene as an important regulator of vascular cambial activity and secondary xylem formation. AHL15 is part of a plant-specific protein family, containing a single AT-hook DNA binding motif and a Plant and Prokaryote Conserved (PPC) domain (Fujimoto et al., 2004; Zhao et al., 2013). *AHL15* homologs have been implicated in several aspects of plant growth and development in Arabidopsis, including flowering time and hypocotyl growth (Street et al., 2008; Xiao et al., 2009), flower development (Ng et al., 2009), vascular tissue differentiation (Zhou et al., 2013), and gibberellin biosynthesis (Matsushita et al., 2007). In addition, we have recently shown that *AHL15* and family members play key roles in plant embryogenesis (Karami et al., 2020b) and –longevity (Karami et al., 2020a). Apart from promoting plant longevity, we also discovered that *AHL15* enhances secondary growth. A more detailed analysis indicated that *AHL15* is a central regulator of vascular cambium activity in *Arabidopsis thaliana* (Arabidopsis) that links the action of the upstream flowering genes *SOC1* and *FUL* to the downstream cytokinin biosynthesis genes *IPT3*, *IPT7* and *LOG4*.

## Results

### *AHL15* promotes secondary growth in Arabidopsis inflorescence stems

In contrast to woody plants that already produce a ring of xylem in young inflorescence stems, herbaceous plants, such as Arabidopsis, produce only a limited amount of xylem without forming a xylem ring in young stems. Secondary growth does occur in older Arabidopsis inflorescence stems and is generally quantified by the number of secondary xylem cell files produced by cambial cell divisions in the interfascicular part of the stem (the area between two bundles) (Nieminen et al., 2015; Fischer et al., 2019). In the Arabidopsis *soc1 ful* double loss-of-function mutant, however, xylem formation is significantly enhanced, resulting in the formation of a xylem ring even in young inflorescence stems (Melzer et al., 2008). In addition, *soc1 ful* mutant plants show extended longevity by the maintenance of vegetative growth from axillary meristems (AMs) (Melzer et al., 2008).

Recently, we have shown that overexpression of the Arabidopsis *AHL15* gene (*p35S:AHL15*) also increases plant longevity (Fig. 1A), similar to *soc1 ful* plants, and that *AHL15* acts downstream of the SOC1 and FUL transcription factors in maintaining AMs in the vegetative phase (Karami et al., 2020a). In view of these data, we analyzed whether *AHL15* overexpression could also promote secondary growth and xylem formation in *p35S:AHL15* inflorescence stems. One-month-old *p35S:AHL15* stems displayed a significant increase in xylem formation compared to wild-type stems (Fig. 1B). In 2- and 3-month-old *p35S:AHL15* stems secondary xylem formation continued, whereas it stopped in 2-month-old wild-type stems (Fig. 1B, C). These results suggested that AHL15 can trigger cambium activity and thereby enhance secondary growth in Arabidopsis inflorescence stems. Arabidopsis inflorescence stems normally produce more secondary xylem than secondary phloem (Altamura et al., 2001).

**Fig. 1.**
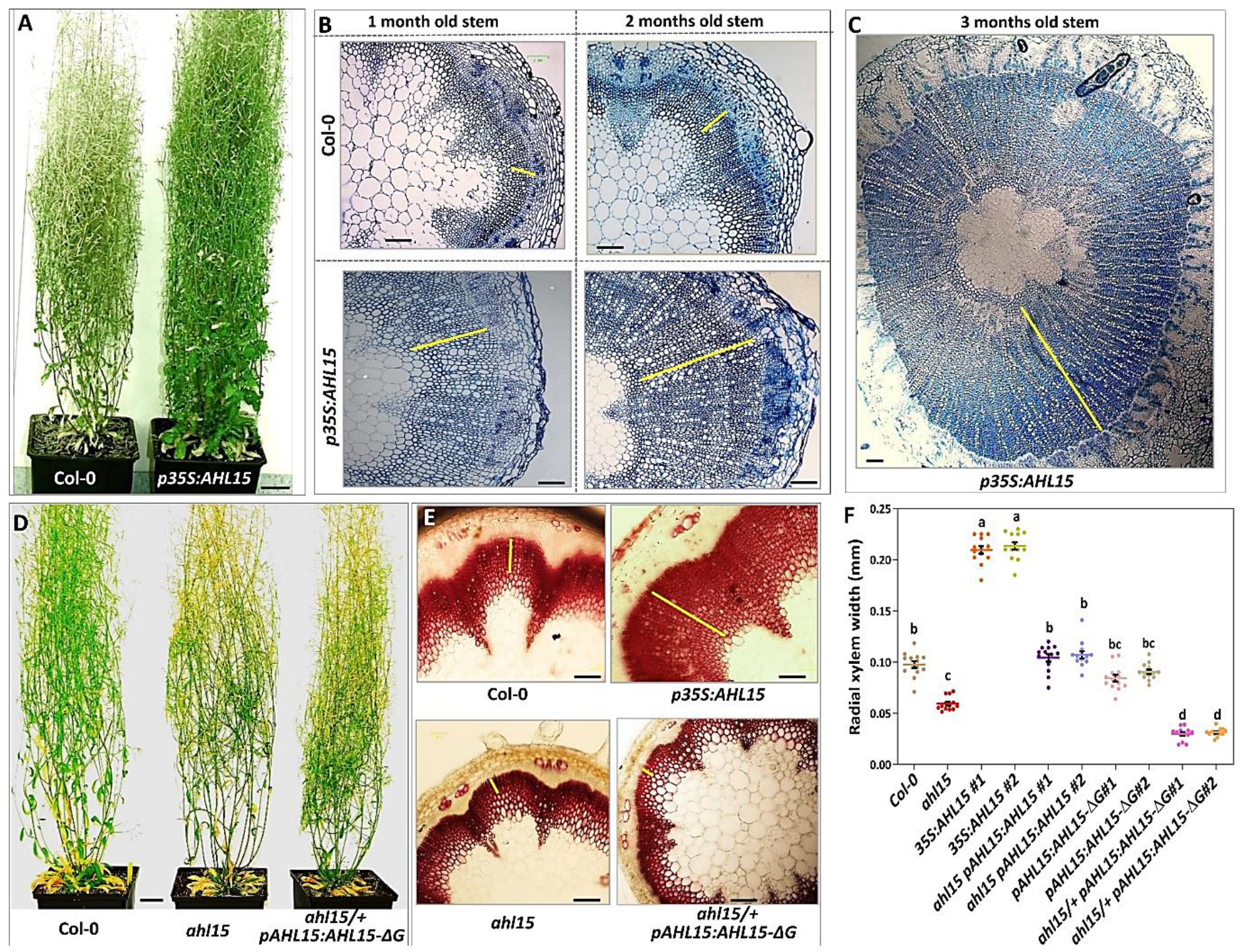
*AHL15* promotes secondary growth in Arabidopsis inflorescence stems. (A) Plant shoot phenotype of a three-month-old wild-type and *p35S:AHL15* plants. (B) Toluidine blue-stained cross-sections of most-bottom base of a one-month-old (left panel) or two-month-old (right panel) inflorescence stem of wild-type and *p35S:AHL15* Arabidopsis plants. (C) Toluidine blue-stained cross-sections of most-bottom base of a three-month-old *p35S:AHL15* inflorescence stem. (D) Shoot phenotypes of two-month-old wild-type (left), *ahl15* (middle) and *ahl15/+ pAHL15:AHL15-ΔG* mutant (right) plants. (E) Phloroglucinol-stained fresh cross-sections of most-bottom base of a one-month-old wild-type (upper panel left), *p35S:AHL15* (upper panel right), *ahl15* (lower panel left), and *ahl15/+ pAHL15:AHL15-ΔG* (lower panel right) inflorescence stems. (F) Quantification of the secondary xylem width (most-bottom base) in interfascicular part of one-month-old wild-type, *p35S:AHL15*, *ahl15* and *pAHL15:AHL15 ahl15,pAHL15:AHL15-ΔG* and *ahl15/+ pAHL15:AHL15-ΔG* main inflorescence stems. Colored dots indicate the average secondary xylem width of 3 random interfascicular region of an individual stem independent plants (n = 9), all the secondary xylem width are presenting the width average per line, horizontal lines indicate the mean and error bars indicate the s.e.m. Different letters indicate statistically significant differences (P < 0.01) as determined by a one-way ANOVA with Tukey’s honest significant difference post hoc test. The yellow bars in (B), (C) and (E) mark the xylem width in the interfascicular cell domain. Scale bars indicate 2 cm in (A) and (D) and 0.06 mm in (B), (C) and (E).

Lignin staining showed that inflorescence stems of one-month-old *p35S:AHL15* plants produced more lignified xylem cells in the interfascicular region than those of wild-type plants (Fig. 1E, F). In contrast, *ahl15* loss-of-function mutant stems contained significantly less lignified xylem cells at interfascicular region than wild-type stems (Fig. 1E, F), even though *ahl15* plants developed and flowered like wild-type plants (Fig. 1D). Introduction of the *pAHL15:AHL15* genomic clone into the *ahl15* mutant background completely restored the secondary xylem growth to wild-type levels (Fig. 1E, F), indicating that the reduction in the xylem cell number was caused by *ahl15* loss-of-function.

Previous studies have shown that expression of a mutant AHL protein (AHL15-ΔG), from which the conserved six amino-acid sequences (GRFEIL) in the PPC domain was deleted, in the heterozygous *ahl15/+* mutant background leads to a dominant-negative effect that overcomes the functional redundancy between the different *AHL* family members (Zhao et al., 2013; Karami et al., 2020a). Whereas *pAHL15:AHL15-ΔG* plants showed wild-type development, as previously reported (Karami et al., 2020a), *ahl15/+ pAHL15:AHL15-ΔG* plants developed even less secondary xylem (Fig. 1D) compared to *ahl15* plants (Fig. 1E, F). The diameter of *ahl15* stems was already smaller than wild-type stems, but *ahl15/+ pAHL15:AHL15-ΔG* stems remained even smaller in size (Supplementary Figure 1). Except for the difference in cambium activity, mutant stems showed the same structure and tissue organization as wild-type stems (Supplementary Figure 2). These results show that *AHL15* plays an important role in increasing cambium activity and secondary growth in Arabidopsis inflorescence stems, and that like with developmental processes such as embryogenesis and AM maturation (Karami et al., 2020a, 2020b), it acts partially redundant with other *AHL* genes.

### SOC1/FUL-repressed cambium-specific *AHL15* expression regulates secondary growth

The above results suggested that *AHL15* is expressed in the vascular cambium area where it regulates cambial cell division activity. Histochemical staining of plants having the *AHL15* promoter β-glucuronidase (GUS) reporter (*pAHL15:GUS*) in wild-type background indicated that *AHL15* is expressed in almost all tissues in 1-week-old inflorescence stems (1-week after bolting) (Fig. 2A), but that the expression became progressively restricted to the vascular and interfascicular cambium zone during stem maturation (Fig. 2A). Interestingly, *AHL15* expression was completely absent from the secondary xylem, whereas it remained expressed in the primary xylem and xylem fibers of the inflorescence stem (Fig. 2A).

**Fig. 2.**
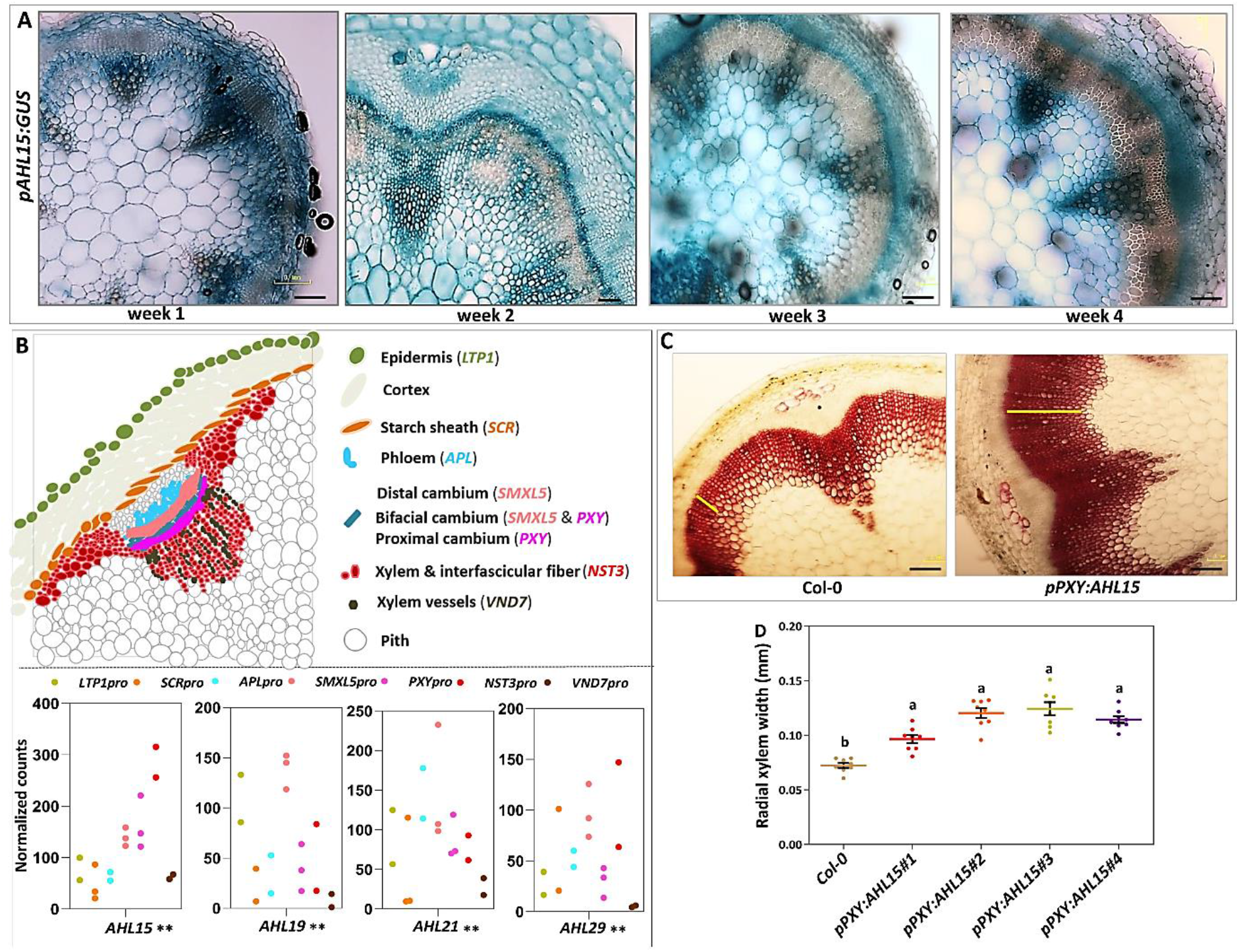
Along with unique expression pattern of*AHL15* in cambium area with other*AHLs*, Cambium-specific *AHL15* expression promotes secondary xylem formation. (A) Cross-sections of most-bottom base of a one-,two-, three- and four-week-old inflorescence stems presenting the expression pattern the *pAHL15:GUS* reporter line (promoter activity). (B) Schematic representation of part of a cross section of an inflorescence stem, including the vascular bundle and the interfascicular regions, in which different stem tissues and the tissue-specific expression domains of genes used for stem transcriptome profiling are indicated (*NST3* (fibers), *VND7*(xylem vessels), *PXY* (proximal cambium), *SMXL5* (distal cambium), *APL* (phloem), *SCR*(starch sheath), *LTP*1(epidermis cells) (upper panel)). Co-expression of *AHL15*, *AHL19, AHL21* and *AHL29* genes at cambium and bundle domains. (lower panel). Normalized read counts of the *AHL* genes in the tissue-specific gene expression atlas generated by combining fluorescence-activated nucleus sorting (FANS) using the promoter-H4-GFP fusions of the indicated marker genes (*NST3*, *VND7*, *PXY*, *SMXL5*, *APL, SCR* and *LTP1*) and combining FANS and laser-capture microdissection with next generation RNA sequencing (Shi et al., 2020). Colored dots indicate the values of two or three biological replicates of RNA isolation obtain by FANS, Normalized gene read counts of the indicated genes among seven different tissues displayed for each replicate individually. *, ** indicates p < 0.05 and 0.01, respectively in LRT (Shi et al., 2020). (C) Phloroglucinol-stained fresh cross-sections of most-bottom base of a one-month-old wild-type (left) and *pPXY:AHL15* (right) inflorescence stems. The yellow bars mark the xylem width in the interfascicular cell domain. (D) Quantification of the secondary xylem width (most-bottom base) in interfascicular part of one-month-old wild-type and *pPXY:AHL15* main inflorescence stems. (5 individual lines). Colored dots indicate the average secondary xylem width of 3 random interfascicular region of an individual stem independent plants (n = 7), all the secondary xylem width are presenting the width average per line, horizontal lines indicate the mean and error bars indicate the s.e.m. Different letters indicate statistically significant differences (P < 0.01) as determined by a one-way ANOVA with Tukey’s honest significant difference post hoc test. Scale bars in (A) and (C) indicate 0.06 mm.

In order to confirm that *AHL15* is expressed in the vascular cambium, we extracted the expression pattern of clade-A *AHL* genes from tissue-specific gene expression atlas generated from different Arabidopsis stem tissues (e.g., xylem vessels, fibers, the proximal and the distal cambium, phloem, phloem cap, pith, starch sheath, and epidermis cells; see Fig. 2B) by combining fluorescence-activated nucleus sorting and laser-capture microdissection with next-generation RNA sequencing (Shi et al., 2020). The extracted *AHL15* expression from this data belonging to the second bottom-most internode of Arabidopsis stem (about 2 weeks after bolting) (Fig. 2B) showed a high level of similarity with the activity pattern of the *pAHL15:GUS* reporter in 2-week-old inflorescence stem up to 4-week old stems. The expression of *AHL15* in the *PXY* and *SUPPRESSOR OF MAX2 1-LIKE PROTEIN 5* (*SMXL5*) cambium sub-domains, the areas where cell division is taking place (Shi et al., 2019), further supported the role of *AHL15* in promoting vascular cambium activity. In line with the functional redundancy between *AHL* genes in promoting cambium activity, we also found *AHL19*, *AHL20*, and *AHL29* to be expressed in the *PXY* and *SMXL5* cambium sub-domains (Fig. 2B).

Altogether, the seemingly overlapping expression of *AHL* genes in the stem cambium zone supports their role in the regulation of cambium activity. To establish whether *AHL* expression in the Arabidopsis vascular cambium is rate-limiting for secondary growth, we expressed *AHL15* under control of the cambium-specific *PXY* promotor (*pPXY:AHL15*) (Fisher and Turner, 2007). Lignin stained one-month-old *pPXY:AHL15* stems showed a significant increase in secondary xylem development compared to wild-type stems (Fig.2 C, D). These results indicate that in wild-type Arabidopsis, secondary growth and xylem development are limited by *AHL* expression, and that cambium-specific enhancement of *AHL15* expression is sufficient to promote cambium activity. Apart from the enhanced cambium activity, *pPXY:AHL15* plants developed and flowered like wild-type plants (Supplementary Figure 3), confirming that the effect of locally enhanced *AHL15* expression on cambium activity is direct, and not caused by changes in developmental timing.

Previously, we have shown that *AHL15* expression is repressed by binding of the SOC1 and FUL MADS-box transcription factors to its up- and downstream genomic regions, and that alleviation of this repression in the *soc1 ful* double mutant leads to delayed maturation of AMs, resulting in the mutant aerial rosette phenotype (Karami et al., 2020a). Comparison of *soc1 ful* double mutant with *soc1 ful ahl15* triple mutant stems showed that the high secondary xylem production in *soc1 ful* stems was dependent on the presence of a functional *AHL15* gene. The interfascicular secondary xylem formation was reduced to wild-type levels in the *soc1 ful ahl15* triple mutant (Fig. 3A, B). This is in line with the previously observed enhanced expression of *AHL15* in *soc1 ful* mutant AMs (Karami et al., 2020a). The co-expression of *AHL15* with *SOC1* and *FUL* in the *PXY* and *SMXL5* cambium sub-domains (Fig. 3C) further supports the negative regulation of *AHL15* by *SOC1* and *FUL* in cambium related domains, which limits secondary growth in the herbaceous wild-type Arabidopsis.

**Fig. 3.**
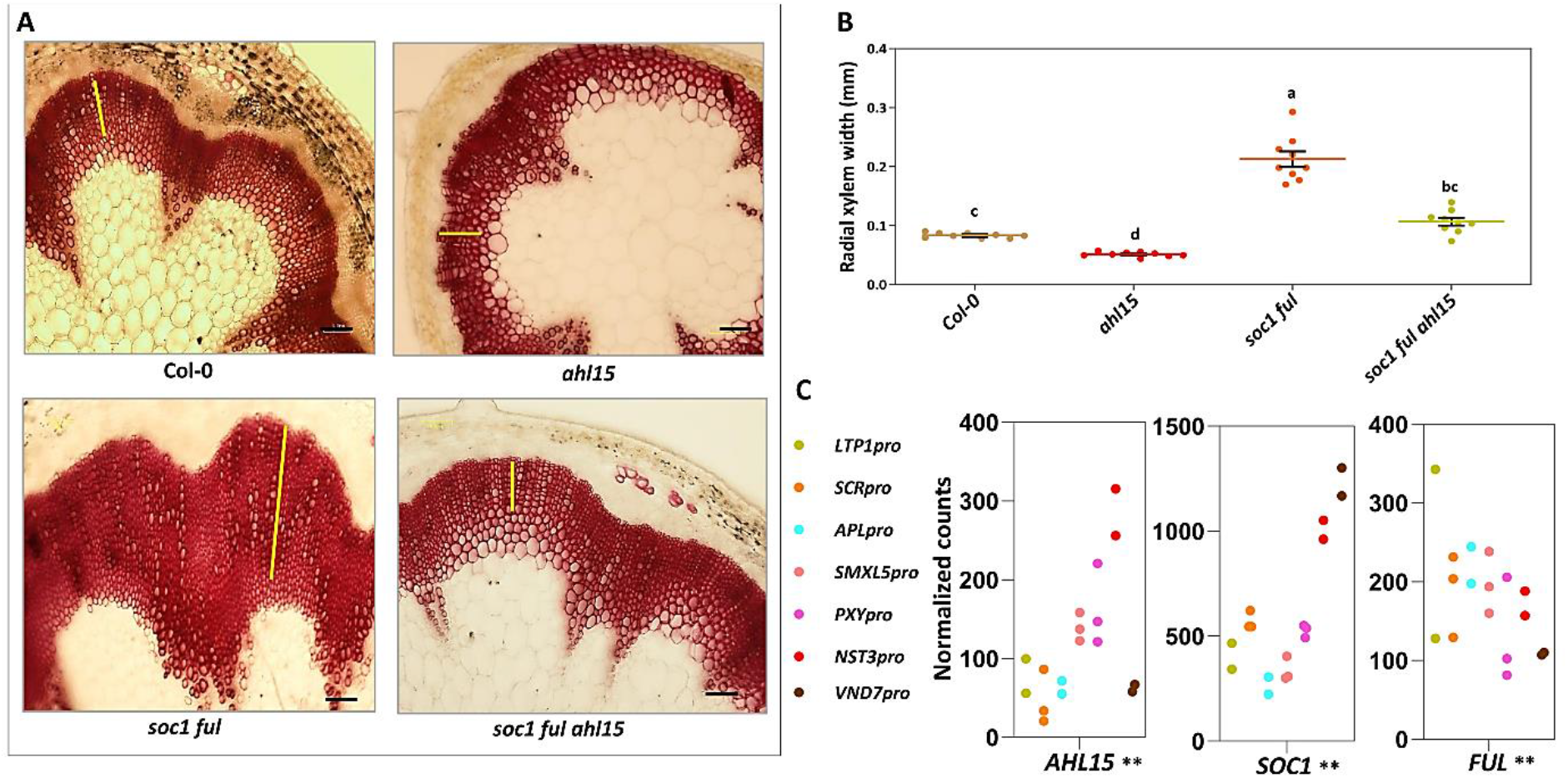
*AHL15* is required for enhanced secondary xylem formation in the *soc1 ful* double mutant. (A) Phloroglucinol-stained fresh cross-sections of most-bottom base of a one-month-old wild-type (upper panel, left), *ahl15* (upper panel, right), *soc1ful* (lower panel, left) and *soc1ful ahl15* (lower panel, right) inflorescence stems. The yellow bars mark the xylem width in the interfascicular cell domain. (B) Quantification of the secondary xylem width (most-bottom base) in interfascicular part of one-month-old wild-type, *ahl15*, *soc1ful* and *soc1ful ahl15* main inflorescence stems. Colored dots indicate the average secondary xylem width of 3 random interfascicular region of an individual stem independent plants (n = 9), all the secondary xylem width are presenting the width average per line, horizontal lines indicate the mean and error bars indicate the s.e.m. Different letters indicate statistically significant differences (P < 0.01) as determined by a one-way ANOVA with Tukey’s honest significant difference post hoc test. (C) Co-expression of *AHL15*, *SOC1* and *FUL* genes at cambium and bundle domains. Colored dots indicate the values of two or three biological replicates of RNA isolation obtain by FANS. Normalized gene read counts of the indicated genes among seven different tissues displayed for each replicate individually. *, ** indicates p < 0.05 and 0.01, respectively in LRT (Shi et al., 2020). Scale bars indicate 0.06 mm.

### *AHL15* promotes secondary growth by increasing cytokinin biosynthesis

In Arabidopsis inflorescence stems, secondary growth is initiated by the occurrence of interfascicular cambium, leading to the formation of a cambium ring. The initial cell divisions marking the starting point of interfascicular cambium formation can be readily observed in one-week-old wild-type stems, but are absent in one-week-old *ahl15* loss-of-function stems (a noticeable feature of this mutant), despite a normal organization of the mutant stem tissues (Fig. 4A). In contrast, one-week-old *p35S:AHL15* inflorescence stems already showed a cambium ring consisting of several layers of cells in the interfascicular region (Fig. 4A). These results indicate an important role for *AHL15* in cambium initiation and the promotion of cambial cell divisions. Previous studies have highlighted that the cell division rate in the procambium and cambium is regulated by the plant hormone cytokinin (Matsumoto-Kitano et al., 2008; Immanen et al., 2016). To test the involvement of cytokinin in the AHL15-enhanced cell divisions, we compared the expression of cytokinin response reporter *pTCS:GFP* (Zürcher et al., 2013) in wild-type, *ahl15* or *p35S:AHL15* inflorescence stems during cambium development. In very young (2-day-old) stems, only a weak *pTCS:GFP* expression could be detected in the procambium zone of stems of all three genetic backgrounds, indicating that *AHL15* does not modulate the cytokinin response in these very young stems stage (Fig.4B, left images). However, three days later, when *pTCS:GFP* signals started to appear in the interfascicular regions of wild-type stems, *pTCS:GFP* expression was still limited to the procambium in *ahl15* stems, whereas *p35S:AHL15* stems displayed a ring of *pTCS:GFP* signal colocalizing with the cambium ring already present in these stems (Fig. 4B, middle images). These overlapping rings of cambium and *pTCS:GFP* expression also became visible in 10-day-old wild-type stems, and were even stronger in *p35S:AHL15* stems (Fig.4B, right images). In 10-day-old *ahl15* stems, however, *pTCS:GFP* expression remained restricted to the cambium in the vascular bundles (Fig.4B, right images). These results suggested that AHL15 might regulate cambium initiation and promote the cambial cell division rate in Arabidopsis stems by altering cytokinin biosynthesis or -response.

**Fig. 4.**
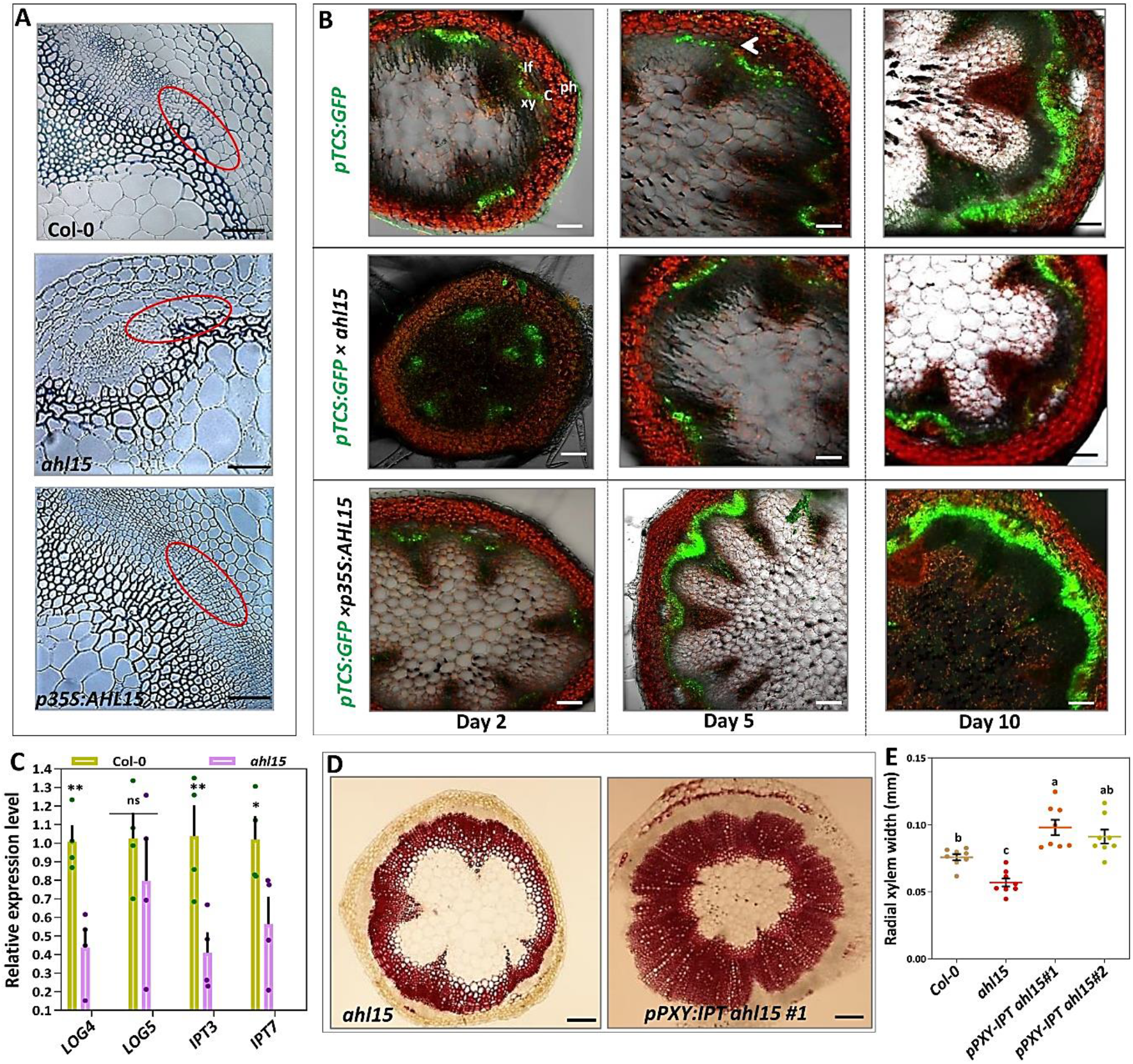
Cytokinin acts downstream of *AHL15* to promote cambium activity. (A) Toluidine blue-stained cross-sections of most-bottom base of one-week-old wild-type (upper), *ahl15* (middle) and *p35S:AHL15* (lower) inflorescence stems. The red circle marks the cambium cell division area in the interfascicular region which is absent in *ahl15* mutant. (B) Confocal images showing the expression of the *pTCS:GFP* cytokinin response reporter cross-sections of most-bottom base of two-, five- and ten-day-old wild-type (upper panel)(arrow indicate the cambium cell division start point in wild-type), *ahl15* (middle panel) and *p35S:AHL15* (lower panel) inflorescence stems. (C) Relative expression of the cytokinin biosynthesis genes *LOG4*, *LOG5*, *IPT3* and *IPT7* by qPCR analysis in the base of ten-day-old wild-type and *ahl15* inflorescence stems. Dots indicate the values of three biological replicates per plant line, bars indicate the mean and error bars indicate the s.e.m. Asterisks indicate significant differences between wild-type and *ahl15* plants (*P < 0.05, **P < 0.01, ns (not significant)), as determined by a two-sided Student’s *t*-test. (D) Phloroglucinol-stained fresh cross-sections of most-bottom base of one-month-old *ahl15* (left) and *ahl15 pPXY:IPT* (right) inflorescence stems. (E) Quantification of the secondary xylem width (most-bottom base) in interfascicular part of one-month-old wild-type, *ahl15* and *ahl15 pPXY:IPT* (2 independent lines) main inflorescence stems. Colored dots indicate the average secondary xylem width of 3 random interfascicular region of an individual stem independent plants (n = 9), all the secondary xylem width are presenting the width average per line, horizontal lines indicate the mean and error bars indicate the s.e.m. Different letters indicate statistically significant differences (P < 0.01) as determined by a one-way ANOVA with Tukey’s honest significant difference post hoc test. Scale bars indicate 0.05mm in A,B and 0.15 mm in D.

To analyze the effect of *AHL15* on cytokinin biosynthesis, the expression of the cytokinin biosynthesis genes *IPT7*, *IPT3*, *LONELY GUY4* (*LOG4*) and *LOG5* (Matsumoto-Kitano et al., 2008) was compared between wild-type and *ahl15* mutant stems. Our qPCR experiments showed that the expression of *IPT3, IPT7*, and *LOG4* was significantly reduced in *ahl15* stems (Fig. 4C), suggesting that *AHL15* acts by activating these cytokinin biosynthesis genes in the cambium area. This was further confirmed by the co-expression of *AHL15*, *IPT3*, *IPT7*, *LOG4*, and *LOG5* in the *PXY* and *SMXL5* cambium sub-domains (Supplementary Fig. 4).

To verify that the reduced secondary xylem formation in *ahl15* mutant stems was caused by a reduced cytokinin biosynthesis, we introduced the *Agrobacterium tumefaciens IPT* gene under the control of the cambium-specific *PXY* promoter (*pPXY:IPT*) in the *ahl15* mutant background (Robson et al., 2004; Van Der Graaff et al., 2001). The presence of the *pPXY:IPT* construct significantly increased the width of interfascicular lignified xylem in *ahl15* stems, bringing it back to wild-type or even higher levels (Fig. 4D-E). Our data demonstrate that local expression of *AHL15* assisted by that of redundantly acting *AHL* genes determines the initiation and activity of interfascicular cambium and thereby the secondary growth of Arabidopsis inflorescence stems by promoting cytokinin biosynthesis.

### *AHL15* promotes secondary growth independent of the PXY-WOX pathway

In Arabidopsis inflorescence stems, the *WUSCHEL HOMEOBOX RELATED 4 (WOX4)* and *WOX14* genes have been shown to promote the rate of cell division in the vascular cambium (Denis et al., 2017; Etchells et al., 2013; Campbell et al., 2016). Inflorescence stems from the *wox4 wox14* double mutant develop significantly less secondary xylem in the interfascicular regions (Etchells et al., 2013). In view of the co-expression of *WOX4*, *WOX14*, and *AHL15* in the *PXY* and *SMXL*5 cambium sub-domains (Fig. 5A) and the fact that loss-of-function mutant inflorescence stems show similar defects in cambium development, we investigated the genetic interaction between *AHL15* and *WOX4/WOX14* in controlling secondary xylem growth. Introduction of *pPXY:AHL15* into the *wox4 wox14 pxy* mutant background did not lead to enhanced secondary growth (Fig. 5B, C). This result indicated that *AHL15* is not responsible for promoting cell division downstream of *WOX4* and *WOX14*, but instead that the PXY-WOX4/14 pathway is required for AHL15 action. However, the *pWOX4:GFP* reporter, which normally labels the vascular bundles and the cambium cells in wild-type stems (Suer et al., 2011), did not show a clear difference in *ahl15* mutant inflorescences stems, indicating that the *WOX* genes are not downstream of AHL15 (Fig. 5 D). Based on these experiments, we conclude that PXY-WOX and AHL15-IPT are parallel pathways that are mutually required to promote cell division in the interfascicular cambium of Arabidopsis inflorescence stems.

**Fig. 5.**
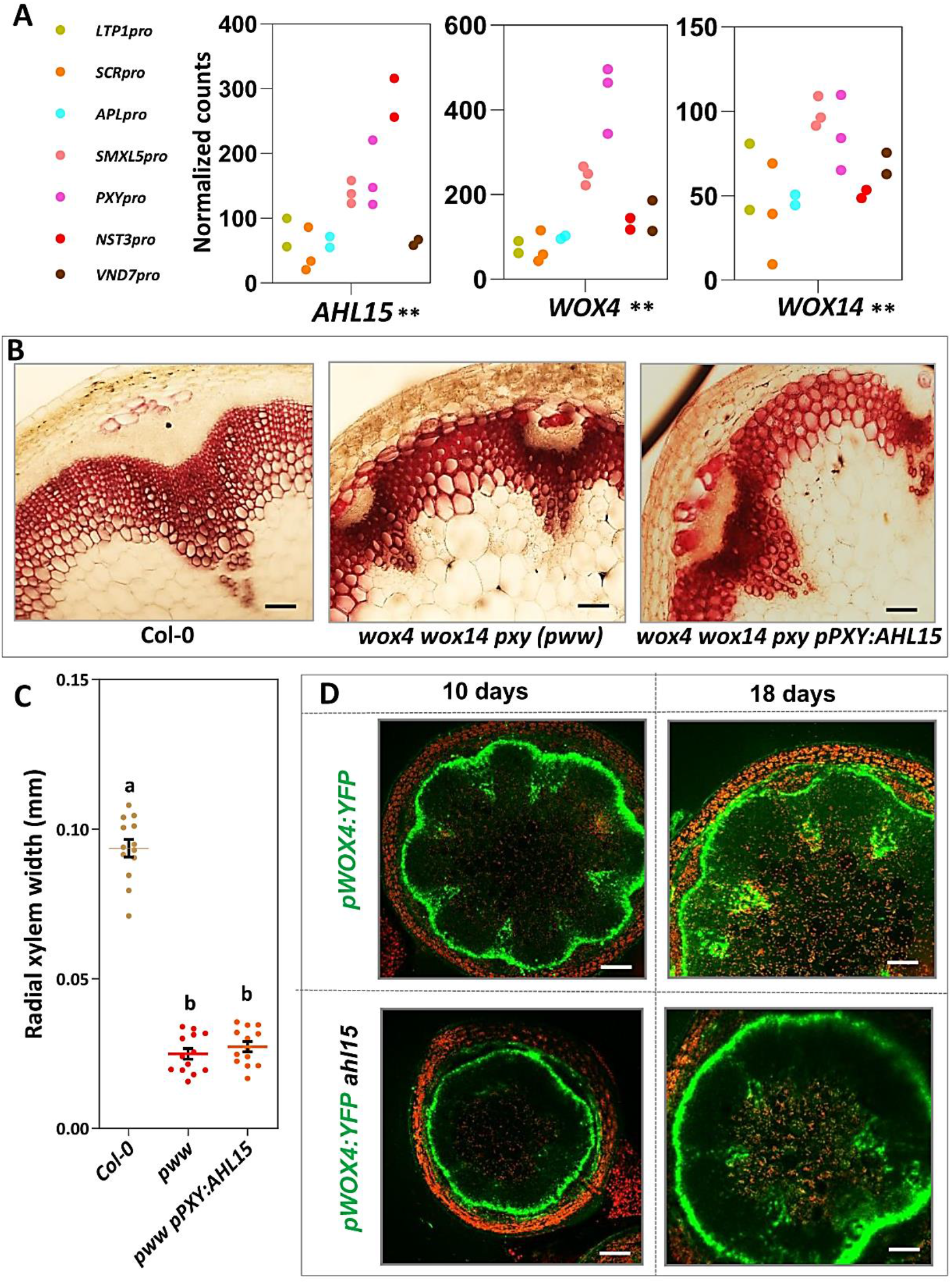
*AHL15-*cytokinin and *PXY-WOX4,14* represent two parallel pathways regulating cambium activity. (A) Co-expression *AHL15*, *WOX4 and WOX14* at cambium and bundle domains. Colored dots indicate the values of two or three biological replicates of RNA isolation obtain by FANS. Normalized gene read counts of the indicated genes among seven different tissues displayed for each replicate individually. *, ** indicates p < 0.05 and 0.01, respectively in LRT (Shi et al., 2020). (B) Phloroglucinol-stained fresh cross-sections of most-bottom base of one-month-old wild-type (left) and *pxy wox4 wox14* (middle) and *pPXY:AHL15 pxy wox4 wox14* (right) inflorescence stems. (C) Quantification of the secondary xylem width (most-bottom base) in interfascicular part of one-month-old wild-type, *pxy wox4 wox14* (*pww*) and *pPXY:AHL15 pxy wox4 wox14* main inflorescence stems. Colored dots indicate the average secondary xylem width of 3 random interfascicular region of an individual stem independent plants (n = 12), all the secondary xylem width are presenting the width average per line, horizontal lines indicate the mean and error bars indicate the s.e.m. Different letters indicate statistically significant differences (P < 0.01) as determined by a one-way ANOVA with Tukey’s honest significant difference post hoc test. (D) Confocal analyses shows activity of *pWOX4-GFP* at 10 and 18-day-old wild-type (up panel) and *ahl15* (low panel) stem. Scale bars indicates 0.06 mm in B and 0.09 mm.

## Discussion

Understanding the molecular mechanisms regulating the activity and functioning of vascular cambium is fundamentally important because the woody biomass in plants derives from the activity of the vascular cambium. The cell division rate of the stem cells of the vascular cambium plays a crucial role in secondary growth: how active the stem cells are in the cambial zone can determine the amount of secondary growth or wood formation (Campbell et al., 2016; Chiang and Greb, 2019). Although previous studies have provided the first insight into the molecular basis regulating cambial activity, our fundamental insight into how initiating and regulated cambial stem cell activity has remained largely unknown.

In this study, we present the role of *AHL15* as a novel regulator in the control of interfascicular cambium activity and secondary xylem formation in Arabidopsis. In particular, in inflorescence stems of *ahl15* loss-of-function mutant plants, we observed a delay in cambium initiation and formation from parenchyma cells and a significant reduction in the amount of secondary xylem formed. By contrast, inflorescence stems of *AHL15* overexpression plants developed an increased amount of secondary xylem. Our data suggest that the *AHL15* gene is a positive regulator of cambial cell proliferation and thereby an important determinant of secondary xylem formation in Arabidopsis inflorescence stems.

*AHL15* belongs to a large family of *AHL* genes in Arabidopsis that have a high degree of functional redundancy among family members (Xiao et al., 2009; Zhao et al., 2013). In line with our recent observations on the redundant action of *AHL15* and other *AHL* genes in controlling axillary meristem maturation (Karami et al., 2020a), and embryogenesis (Karami et al., 2020b), the results presented here suggest that *AHL* genes also act redundantly in regulating interfascicular cambium activity. The overlapping expression of *AHL15*, *AHL19*, *AHL20*, and *AHL28* in the *PXY*/*SMXL5*-marked cambium region of inflorescence stems is in line with their redundant role in controlling cambial cell proliferation. Despite this redundancy, and as observed for axillary meristem maturation (Karami et al., 2020a), *AHL15* seems to play a key role in this process, since secondary growth is significantly reduced in the *ahl15* loss-of-function mutant background.

The cell division in the cambial zone is known to be controlled by both genetic and hormonal factors (Immanen et al., 2016; Oles et al., 2017; Fischer et al., 2019). Based on the current model, the TDIF-PXY-WOX and cytokinin/auxin pathways act in parallel to regulate cell division in the vascular cambium (Fig. 6) (Fischer et al., 2019). In the first pathway, the TDIF peptide, belonging to the CLE family of peptides, is produced in the phloem and subsequently transferred to the cambium where it binds to the PXY receptor-like kinase. Upon peptide binding, PXY activates the expression of the cambium-specific transcription factors WOX4 and WOX14, which promote cell proliferation in the cambium (Fig. 6). Our results show that AHL15 promotes the timing of cambial cell initiation, proliferation, and maintenance afterward in parallel to and dependent on the TDIF-PXY-WOX pathway by increasing the expression of cytokinin biosynthesis genes (Fig. 6). Interestingly, it has been shown that the cambial cell division and secondary xylem production are significantly increased upon elevated cytokinin levels in the stem of poplar trees (Immanen et al., 2016).

**Fig. 6.**
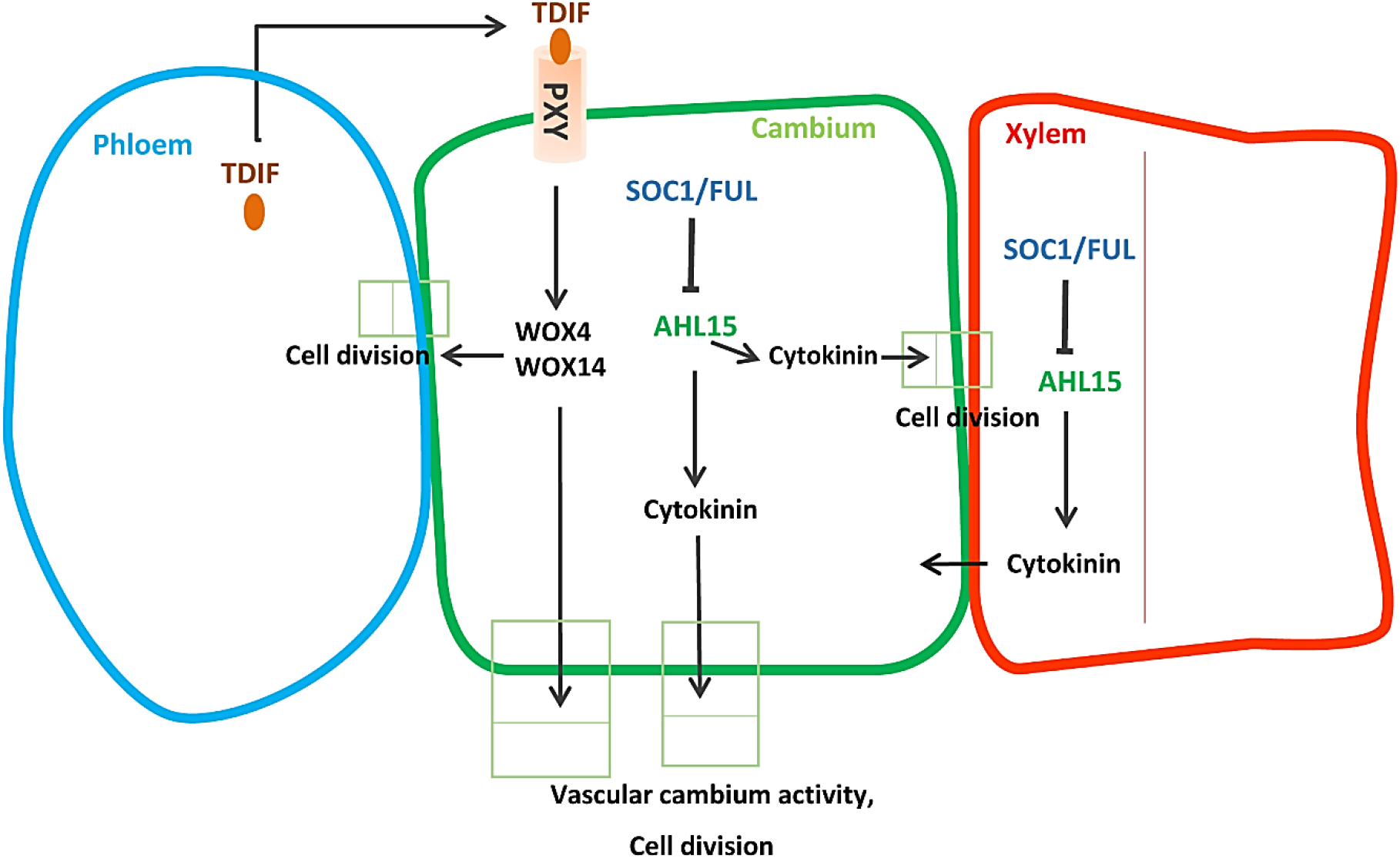
Proposed model for*AHL15* and its key role in promoting cambium activity. Close up of cambium area with different units include phloem (blue), cambium (green), and xylem (red). Peptide TDIF is synthesized in the phloem, travels through the apoplastic space to cambium cells, where it binds to its receptor PXY. PXY subsequently promotes the proliferation of the cambium cells by activating the expression of *WOX4* and *WOX14*. In a parallel pathway, AHL15 promotes cambium activity independent of PXY-WOX and in downstream of SOC1 and FUL and upstream of cytokinin biosynthesis either in cambium cells or in xylem. Blunt-ending lines indicate repression, arrows indicate promotion.

We have previously shown that the MADS-box transcription factors SOC1 and FUL repress AHL15 expression by directly binding to the AHL15 upstream and downstream regions (Karami et al., 2020a). In addition, we show here that the enhanced secondary growth in the *soc1 ful* double mutant inflorescence stems is dependent on the presence of a functional *AHL15* gene, indicating that SOC1 and FUL also act as repressors of secondary growth upstream of *AHL15* (Fig. 6). We conclude that PXY-WOX and SOC1/FUL-AHL15-IPT are parallel pathways that are mutually required to promote cell division in the interfascicular cambium of Arabidopsis inflorescence stems.

It has been reported that AHL3-AHL4 heterodimers control the boundary between the procambium and xylem axis in Arabidopsis roots (Zhou et al., 2013). Single and double loss-of-function mutants of *AHL3* and *AHL4* display ectopic protoxylem vessels and ectopic metaxylem vessels in the procambial region adjacent to the xylem axis (Zhou et al., 2013). Since the cytokinin response is altered in the *ahl4* loss-of-function mutant and similar vascular defects have been observed for cytokinin-defective mutants (Zhou et al., 2013), this confirms the strong relationship between *AHL* genes and cytokinin in cambium initiation and cell division, and in the resulting xylem formation.

How exactly cytokinin promotes cambial cell divisions is still not understood. Cytokinin likely influences the cell cycle of the cambium stem cells. Because cytokinin and auxin usually work together, cytokinin may exert its effect on the cambial cell division by reducing the concentration of auxin in cambium cells by up-regulating the levels of PIN auxin efflux carriers (Duclercq et al.,2009; Šimášková et al., 2015). However, this should be confirmed by future studies.

In conclusion, our studies show that *AHL* genes positively regulate both cambium activity and axillary meristem outgrowth in Arabidopsis, downstream of the SOC1 and FUL transcription factors and upstream of cytokinin biosynthesis. Our findings represent an important step forward in our understanding of the increased vascular cambium activity in *soc1 ful* mutant plants. In particular, our findings show that it is possible to increase secondary growth without affecting the flowering time and fruit and seed development of plants, which will eventually be helpful for plant biomass improvement.

## Methods

### Plant material, growth conditions, and transgenic *Arabidopsis* lines

All Arabidopsis mutant- and transgenic lines used in this study are in the Columbia (Col-0) background. The *ahl15* mutant and the *pAHL15:AHL15*, *pAHL15:GUS*, *ahl15/+ pAHL15:AHL15-ΔG* transgenic lines (Karami et al., 2020a) and the *p35S:AHL15* transgenic line (Karami et al., 2020a) were previously described. The *soc1-6 ful-7* double mutant (Melzer et al., 2008), the *pxy wox4 wox14* mutant and the *pTCS:GFP* (Bruno and Jen, 2008) and *pWOX4:YFP* (Suer et al., 2011) reporter lines were obtained from the Nottingham Arabidopsis Stock Centre (NASC). Seeds were surface sterilized with 40% bleach solution followed by four times washing with sterile water and germinated, after three days incubation at 4°C, on ½ MS (Murashige and Skoog, 1962) medium containing 1% sucrose, 0.7% agar at 21°C and a 16 hours photoperiod. Two-week-old seedlings were transferred to soil and grown at 21°C, 65% relative humidity and a 16 hours (long day: LD) photoperiod, and potted plants were photographed with a Nikon D5300 camera.

### Plasmid construction

To generate the *pPXY:AHL15* construct, upstream regions of approximately 3 kb from the ATG initiation codon of the *PXY (AT5G61480)* gene was amplified from ecotype Columbia (Col-0) genomic DNA using the forward (F) and reverse (R) PCR primers indicated in Supplementary Table 1. The resulting fragments were first recombined into pDONR207 by a BP reaction and subsequently cloned upstream of the *AHL15* cDNA fragment in previously made destination vector *pGW-AHL15* (Karami et al., 2020a) by a LR reaction according to Gateway cloning procedures (Clonetech). To generate the *pPXY:IPT* construct, the *AHL15* gene in the *pPXY:AHL15* construct was replaced by a *Kpn*I and *Xba*I flanked fragment containing the *IPT* gene from *Agrobacterium tumefaciens* (Van Der Graaff et al., 2001). All binary vectors were introduced into *Agrobacterium tumefaciens* strain AGL1 by electroporation (den Dulk-Ras and Hooykaas, 1995). Subsequently, Arabidopsis Col-0, *ahl15*, and *wox4wox14pxy* plants were transformed using the floral dip method (Clough and Bent, 1998).

### RNA preparation and quantitative real-time PCR (qRT-PCR)

RNA was isolated from rosette base nodes, and the basal part of inflorescence stems (about 0.5 cm above rosette base) using the RNEasy© kit (Qiagen). First-strand cDNA was synthesized using the RevertAid RT Reverse Transcription kit (Thermo Fischer Scientific). Quantitative PCR was performed on three biological replicates along with three technical replicates using the SYBR-green dye premixed master-mix (Thermo Fischer Scientific) in a C1000 Touch© thermal cycler (BIO-RAD). CT values were obtained using Bio-Rad CFX manager 3.1. The relative expression level of genes was calculated according to the 2^-ΔΔCt^ method (Livak and Schmittgen, 2001), and expression was normalized by using the *β-TUBULIN-6* gene as a reference, analyzed and plotted into graphs in GraphPad Prism 8. The primers used for each gene are described in Supplementary Table 1.

### Histology and microscopy

To analyze stem secondary growth and lignification, the base part (bottom-most) of the main inflorescence stem was cut and sectioned freshly using a razor blade (Wilkinson Sword) (very thin-fresh free-hand cross-sections), which were kept on the ice up to the staining in 3% phloroglucinol-HCL (0.3 g phloroglucinol, 10 ml absolute ethanol, 5 ml 37 % HCL), using a protocol from Mitra & Loqué 2014 (Pradhan Mitra and Loqué, 2014). Sections were further analyzed under a light microscope (Nikon eclipse Ci-E/Ci-L). The secondary xylem (SX) width was measured with ImageJ in the way three randomly selected interfascicular parts of the individual stem were measured and divided by 3 to have SX width per stem. For embedded sections, the basal part (bottom-most) of the 1or 2 -month old main inflorescence stem segments at least 1cm in length (including the stem base) were harvested and were fixed overnight in 4 % formaldehyde in a 50 mM phosphate solution. The stems were dehydrated in a graded ethanol series (70 %, 80 %, 90 %, 96 %, 100 %) and embedded in epoxy resin molds. Sections (3-4μm) cut by rotary microtome (Leica RM2265) were stained with 0.01 % aqueous toluidine blue, before being analyzed under a light microscope (Nikon eclipse Ci-E/Ci-L).

Histochemical β-glucuronidase (GUS) staining was performed as described previously (Anandalakshmi et al.,1998). The stem segments were cut one centimeter above the basal part and placed in the vial tubes containing cold acetone 90% on ice, after vacuum infiltration and keeping them on ice the protocol followed with two-time washing and vacuum infiltration afterward transferring to GUS staining solution (containing X-gluc), at the end staining reaction was allowed to proceed around 2 hours at 37°C in the dark. After rehydration in a graded ethanol series (75, 50, and 25 %), the stained tissues were cut freshly by razor blade (very thin-fresh free-hand cross-sections) and observed and photographed using a light microscope (Nikon eclipse Ci-E/Ci-L). For quantitative analyses, at least seven plants were analyzed for each data point. Cellular and subcellular localization of GFP and YFP protein in fresh hand section stem on microscopy slide covered with coverslip were visualized using the confocal laser scanning microscope (ZEISS-003-18533) with a 534 laser, 488 nm LP excitation and 500-525 nm BP emission filters for GFP and YFP signals. All Photo-based measurements performed in ImageJ then analyzed and plotted into graphs in GraphPad Prism 8.

### Transcriptome profiling of stem tissues

Based on what has been described by Shi et al., 2020, the free-hand cross-sections of the second bottom-most internode of one-month-old Arabidopsis inflorescence stems prepared. Sections belong to different GFP tagged promoter reporter lines (*NST3pro:H4-GFP, VND7pro:H4-GFP, PXYpro:H4-GFP, SMXL5pro:H4-GFP, APLpro:H4-GFP, SCRpro:H4-GFP, LTP1pro:H4-GFP*) corresponding for xylem and interfascicular fiber, xylem vessels, proximal cambium, distal cambium, phloem region, starch sheath and epidermis respectively. GFP positive and negative nucleus confirmed by confocal microscopy, nucleus isolated (15,000 nuclei per sample), and sorted (three replicate for each sample type). After performing RNA isolation, cDNA library prepared for next-generation sequencing. FANS-derived datasets present in this story as an expression of a particular gene in transcriptome data stand for normalized gene read counts of the indicated genes among seven different tissues displayed for each replicate individually. *, ** indicates p < 0.05 and 0.01, respectively in likelihood-ratio test (LRT), null hypothesis to be rejected in the LRT is that among the seven different tissues, genes have similar expression. Read counts analyzed and plotted into graphs in GraphPad Prism 8.

## Acknowledgements

We thank N. Savant and F. van der Klauw, for generating *PXY-AHL15* and *PXY-IPT* constructs respectively, Gerda Lammers and Merijn de Bakker for their help with microscopy. We are grateful to Ward de winter, Jan Vink, Nick Surtel and Mariel Lavrijsen for their supports.

## Author contributions

A.R. designed and performed the majority of Arabidopsis experiments, with contributions from A.L. A.L analyzed the *PXY-AHL15* stem. O.K. and R.O supervised the project. D.S and T.G generated the transcriptome profiling of stem tissues. A.R., O.K. and R.O. wrote the manuscript.

---

**Supplementary Figure 1.**
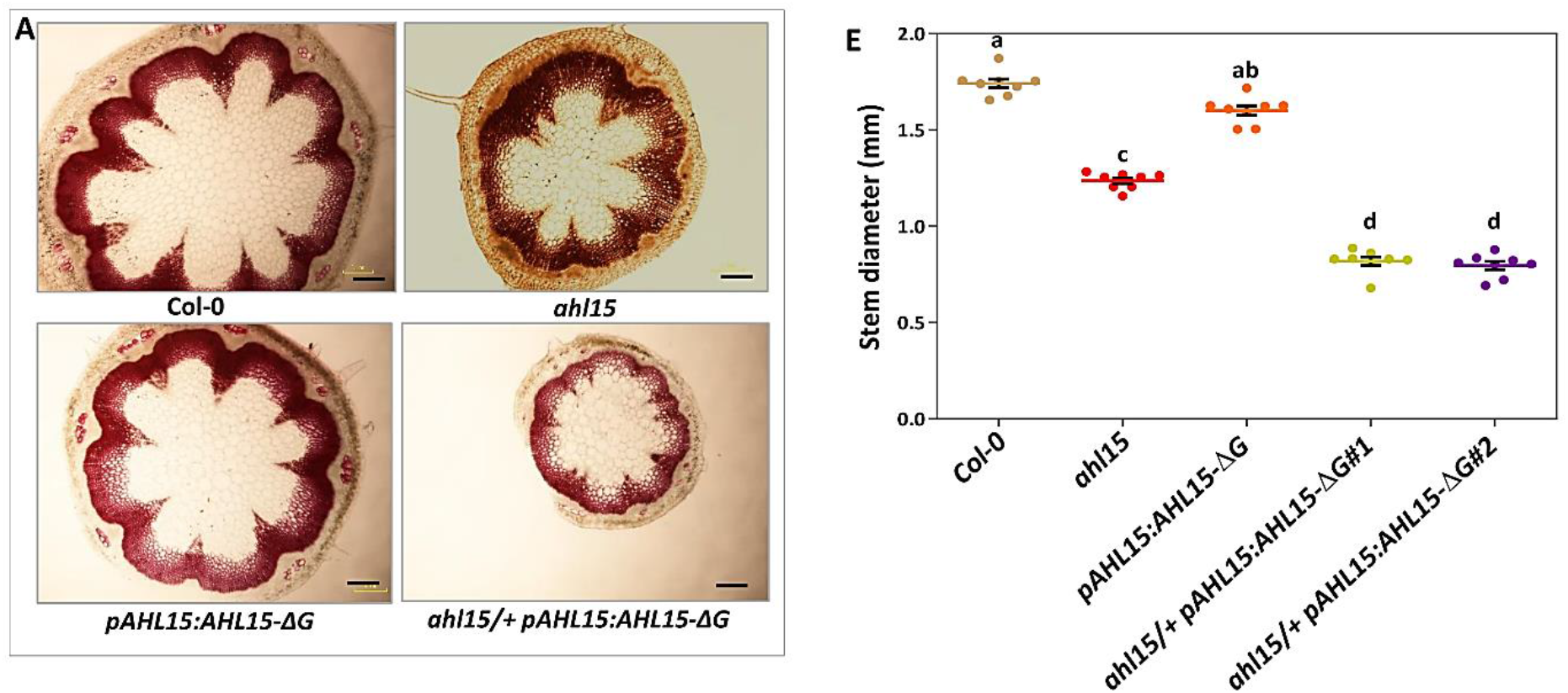
*AHL* genes are essential for secondary growth of Arabidopsis inflorescence stems. (A) Phloroglucinol-stained fresh cross-sections of most-bottom base of one-month-old wild-type (up panel, left), *ahl15* (up panel, right), *pAHL15:AHL15-ΔG* (down panel, left), and *ahl15/+ pAHL15:AHL15-ΔG* (down pane, right) inflorescence stems. (B) Quantification of the stem diameter in one-month-old stem wild-type, *ahl15*, *pAHL15:AHL15-ΔG*, and *ahl15/+ pAHL15:AHL15-ΔG* plants. Colored dots indicate the number of lateral branches per plant (n = 14 (in B) or 10 (in C) biologically independent plants), horizontal lines indicate the mean and error bars indicate the s.e.m. Different letters indicate statistically significant differences (P < 0.01) as determined by a one-way ANOVA with Tukey’s honest significant difference post hoc test. Scale bars indicate 0.15 mm.

**Supplementary Figure 2.**
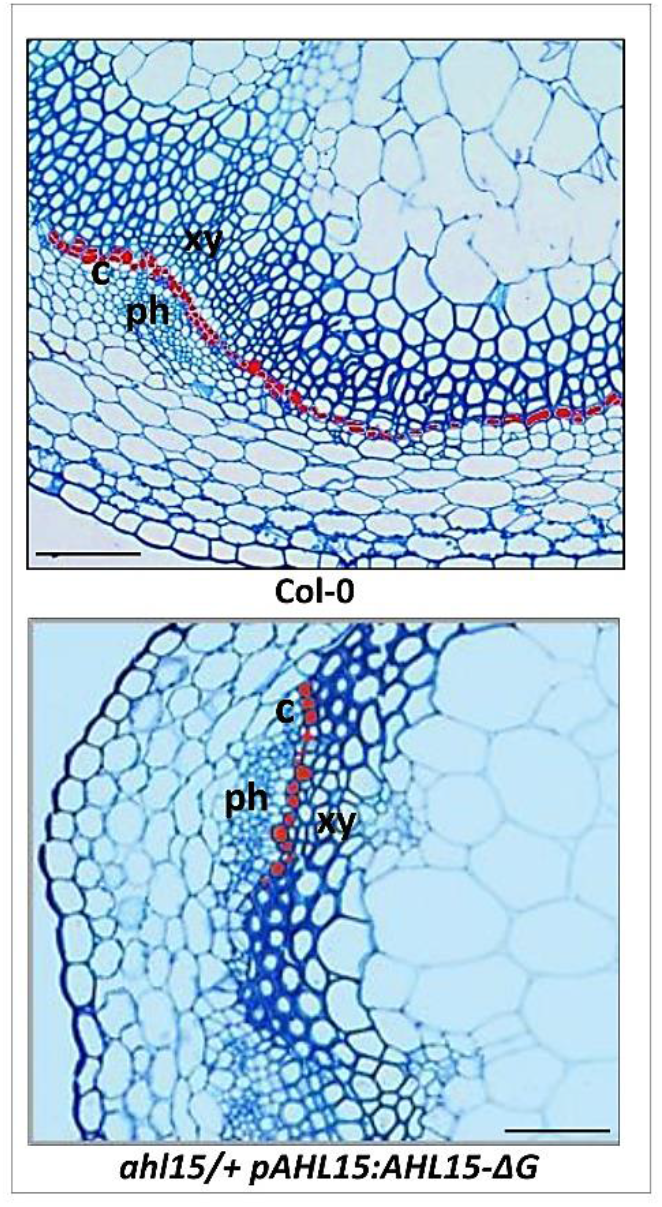
*ahl15/+ pAHL15:AHL15-ΔG* displays normal and organized structure of bundle and interfascicular region compare to wild-type along with delay in cambium initiation. Toluidine blue-stained cross-sections of most-bottom base of two-week-old wild-type (upper) and *ahl15/+ pAHL15:AHL15-ΔG* (lower) inflorescence stems. The red dots mark the cambium/procambial cells in the interfascicular and bundle regions of wild-type and *ahl15/+ pAHL15:AHL15-ΔG* respectively. Scale bars indicate 0.05mm.

**Supplementary Figure 3.**
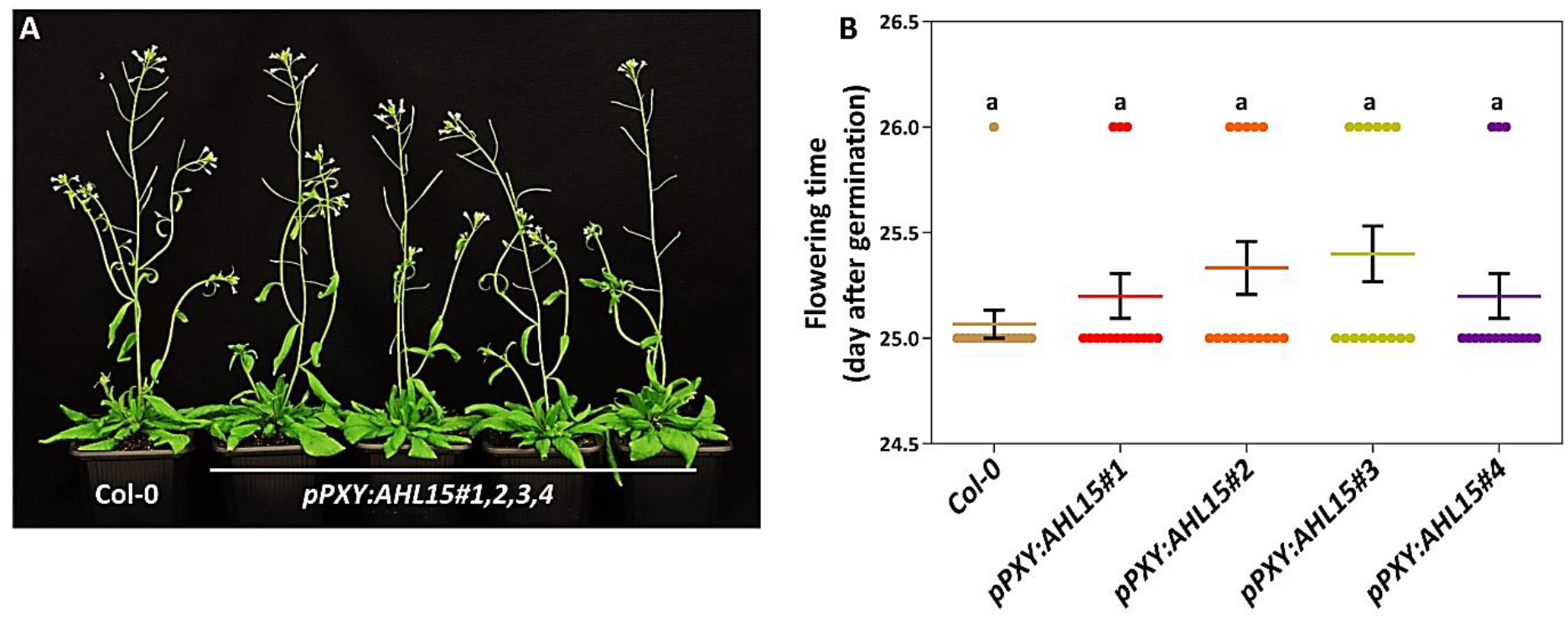
Cambium-specific *AHL15* overexpression does not affect branching or flowering time in Arabidopsis. (A) The shoot phenotype of 5-week-old wild-type and *pPXY:AHL15* (4 independent lines) plants. (B) Quantification of the flowering time of wild-type and *pPXY:AHL15* plants, as presented in (A). Colored dots indicate the flowering time of individual plants in days after germination (n = 15 independent plants per line), horizontal lines indicate the mean and error bars indicate the s.e.m. Different letters indicate statistically significant differences (P < 0.05) as determined by a one-way ANOVA with Tukey’s honest significant difference post hoc test.

**Supplementary Figure 4.**
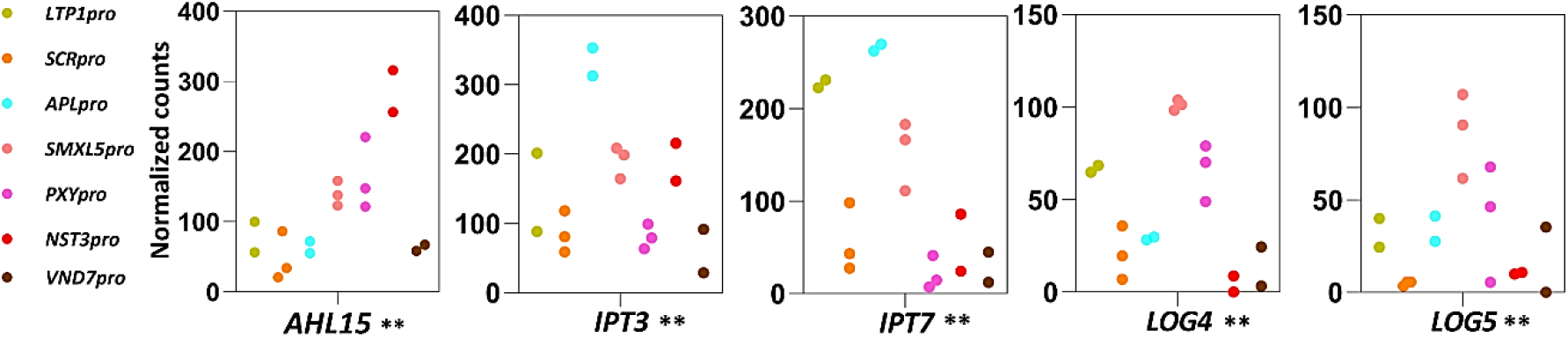
*AHL15* and cytokinin biosynthesis genes are co-expressed in the cambium domain. Co-expression of *AHL15* with *IPT3*, *IPT7, LOG4* and *LOG5* at cambium and bundle domains. Colored dots indicate the values of two or three biological replicates of RNA isolation obtain by FANS. Normalized gene read counts of the indicated genes among seven different tissues displayed for each replicate individually. *, ** indicates p < 0.05 and 0.01, respectively in LRT (Shi et al., 2020).

**Supplementary Table 1.**
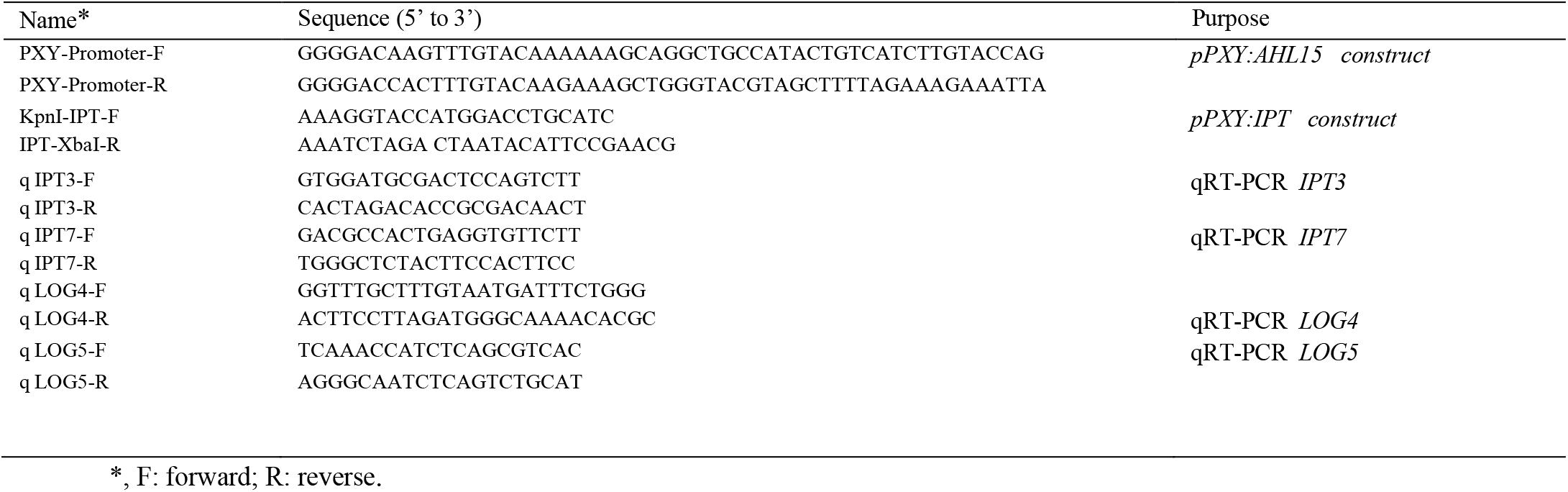
Primers used for cloning, genotyping and qRT-PCR

